# Prediction of Enzyme function using interpretable optimized Ensemble learning framework

**DOI:** 10.1101/2025.01.28.635225

**Authors:** Saikat Dhibar, Sumon Basak, Biman Jana

**Author notes:** Both the authors contributed equally to this work (Saikat Dhibar, Sumon Basak).

## Abstract

Accurate prediction of enzyme function, particularly for newly discovered uncharacterized sequences, is immensely important for modern biology research. Recently machine learning (ML) based methods have shown promises. However, such tools often suffer from complexity in feature extraction, interpretability, and generalization ability. Here we present an interpretable ML method, SOLVE that addresses these issues by using only combination of tokenized subsequences from the protein’s primary sequence for classification. Its optimized ensemble learning framework improves prediction accuracy, distinguishes enzymes from non-enzymes, and predicts enzyme commission (EC) numbers for mono- and multi-functional enzymes. Additionally, SOLVE provides interpretability through Shapley analyses, identifying functional motifs at catalytic and allosteric sites of enzymes. By focusing only on primary sequence data, SOLVE simplifies high-throughput enzyme function prediction for functionally uncharacterized sequences and outperforms existing tools. With high prediction accuracy and its identification ability of functional regions, SOLVE can become a promising tool in different fields of biology and therapeutic drug design.

## 2. Introduction

Enzymes, as biocatalysts, expedite biochemical reactions within cellular frameworks. Their functional categorization has extensive applications in biotechnology^1^, healthcare^2^, and metagenomics^3^. In pharma companies, enzymes facilitate processes such as biosynthesis and polymer recycling.^4^ Many bacterial enzymes in the human gut microbiota require functional annotation, as alterations in bacterial colonies are associated with irritable bowel disease (IBD) and obesity.^5^ Functional annotations would assist medical science by underpinning species that produce the requisite enzymes, aiding disease treatment. To accurately determine an enzyme’s function using biochemical assays, wet labs require significant investments in pricey reagents, extensive experimental time, and the expertise of skilled researchers.^6^ As of May 2024, UniProtKB/Swiss-Prot^7^ contains 283,902 manually annotated enzyme sequences, representing just 0.64% of the total 43.48 million enzyme sequences in the database. Thus, experimental methods become potentially unsustainable in the omics era, when large-scale genome projects continuously add new enzyme sequences to databases. Therefore, computational tools provide valuable guidance for experiments with models that are efficient, cost-effective, reproducible, and maintain high accuracy.

Enzymes are classified using an ontological system known as the Enzyme Commission (EC) number, which organizes them based on the types of reactions they catalyze. This system is structured hierarchically into four levels, which we denote as *L1, L2, L3, and L4*. At the first level (*L1*), enzymes are divided into seven major classes: (i) oxidoreductases, (ii) transferases, (iii) hydrolases, (iv) lyases, (v) isomerases, (vi) ligases, and (vii) translocases. As the classification becomes more specific, the second level (*L2*) designates the subclass, the third level (*L3*) identifies the sub-subclass, and the fourth level (*L4*) specifies the substrate or substrate group upon which the enzyme acts. This tiered system ensures detailed and precise categorization of enzymes, from broad functional roles to specific substrate interactions. *In silico* methods can annotate a novel protein functionally with accurate EC number prediction at any level. Homology-based^8,9^, physicochemical^10,11^, structural^12–14^, and sequence-derived^15–21^ properties have been explored in the last few decades—even specific methods combined multiple properties for function predictions.^16,22,23^

Nonetheless, each of these methods exhibits distinct disadvantages alongside their advantages. For instance, BLAST identified adenylosuccinate lyase (involved in nucleotide biosynthesis), fumarase (involved in the citric acid cycle), and aspartate ammonia lyase (involved in amino acid metabolism) as homologs albeit they perform dissimilar functions.^24^ As of March2024, the Protein Data Bank (PDB) contains 103,972 experimentally determined enzyme structures, representing only a tiny fraction of enzymes catalogued in UniProtKB. Although AlphaFold^25^ has enabled high-throughput structure prediction of proteins, classifying millions of them with structural information is still a computationally intense task. Sequence-based models primarily depend on manually selected intuitive descriptors, often sequence-length dependent. Several methods, like pse-AAC, have been proposed to derive sequence-length-independent descriptors from sequence-length-dependent ones.^11,26^ Nevertheless, these methods usually necessitate manual intervention and may introduce errors through the standardization of dimensionality.

Computational approaches have demonstrated significant potential in elucidating both protein functions and their associated functional landscape.^27–33^ The first known use of machine learning (ML) for enzyme annotation dates back to 1997.^34^ Since then, numerous tools have been developed with improving accuracy.^10,13,15,22,23,35–38^ Several methods have been employed in enzyme function prediction models, including explainable artificial intelligence (XAI) tools like *k*-Nearest Neighbor (*k*NN)^37,39^, Support Vector Machine (SVM)^15,21,40,41^, and more recently, neural networks such as N-to-1 neural networks^20^, Artificial Neural Networks (ANN)^42,43^, Convolutional Neural Networks (CNN)^44,45^, Recurrent Neural Networks (RNN)^23^. These models were trained using six primary enzyme classes (*L1*), and while effective, many were developed prior to the inclusion of translocases as the 7th enzyme class in 2018 ^46^,making them a bit outdated. Models developed after that, likeECPred^47^, ProteInfer^48^, CLEAN^49^, DeepEC^50^, DeepECTransformer^51^, ECPICK^44^ were trained on seven primary classes. Despite the remarkable capabilities exhibited by these tools, the scope of improvement persists in feature extraction, model interpretability, and the adaptation of these methodologies to novel sequence datasets training of ML models with minimal and low sequence similarity threshold datasets. Furthermore, a major limitation is their inability to reliably differentiate between enzyme and non-enzyme sequences, leading to the potential mis-assignment of an EC number to non-enzyme proteins when presented with novel sequences. A large-scale community-based Critical Assessment of protein Function Annotation (CAFA)^52^ revealed that nearly 40% of computational enzyme annotation is erroneous. Henceforth, the community requires novel tools capable of automating feature extraction, employing memory-efficient algorithms, and achieving highly accurate enzyme function predictions to accelerate biomedical research and drug development.

In this study, we have developed an XAI model, SOLVE (Soft-Voting Optimized Learning for Versatile Enzymes), designed to classify novel sequences as enzymes or non-enzymes and further determine whether they are mono- or multifunctional. The overview of the SOLVE method is demonstrated in Figure 1. SOLVE identifies the specific *L1* class for monofunctional enzymes, while for multifunctional enzymes, it predicts the relevant *L1* classes. Our model extends to *L4* classification, providing finer granularity in substrate-binding class prediction. SOLVE operates on features extracted directly from the raw primary sequences of proteins. Unlike traditional approaches that depend on predefined biochemical features of protein sequences, our method captures the full spectrum of sequence variations, allowing the model to learn intricate patterns inherent in the protein sequences. Numerical tokenization enhances computational efficiency by reducing the dimensionality of the input space while preserving critical sequence information. To the best of our knowledge, no other contemporary study has successfully extended from enzyme-nonenzyme binary classification to *L4* substrate binding multilabel multiclass prediction with such excellent to moderate accuracy. This method not only surpasses most existing algorithms in classification accuracy but also offers a more interpretable model by directly linking specific subsequence patterns to enzyme activity, thereby providing novel insights into enzyme structure-function relationships.

**Figure 1:**
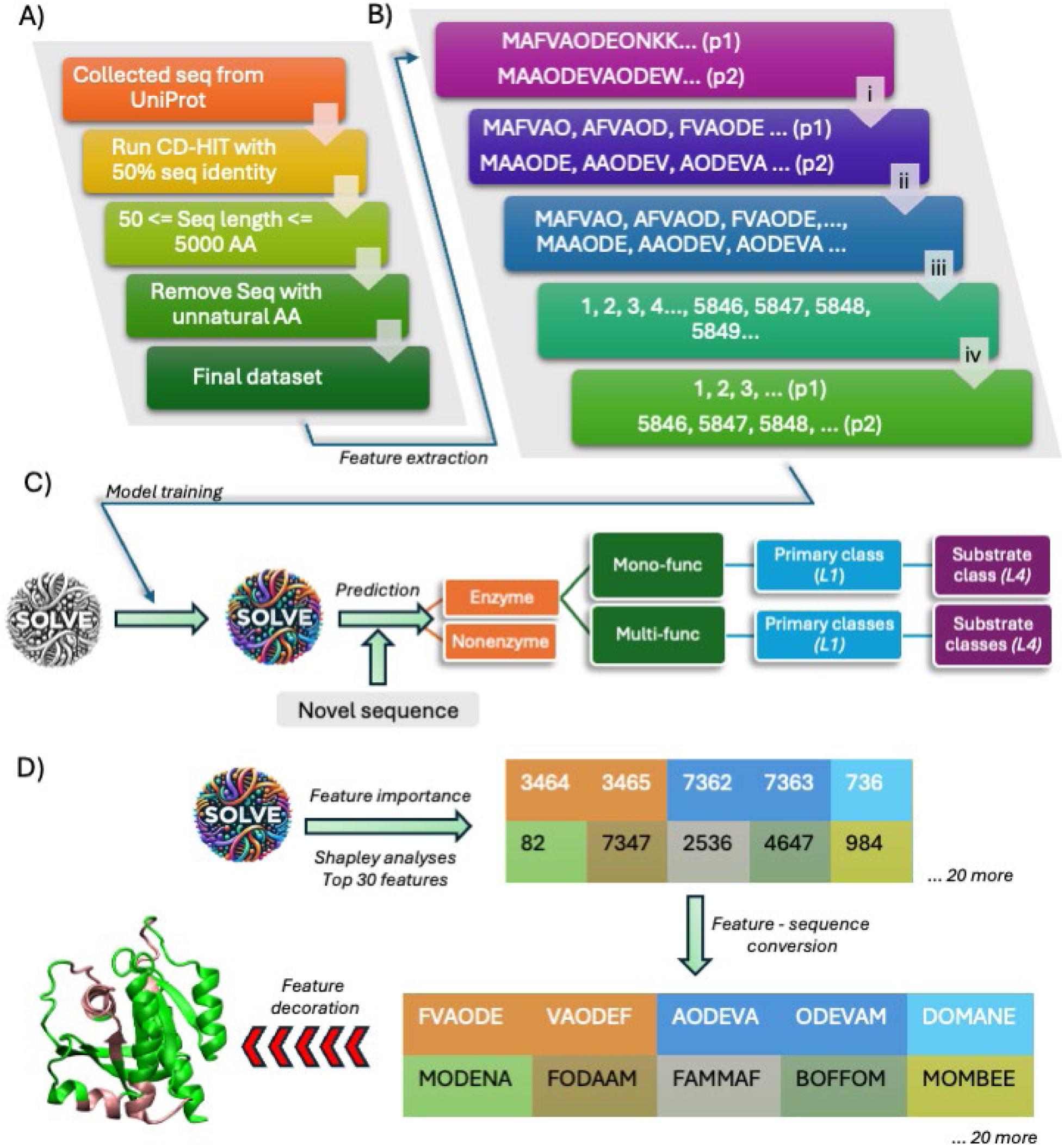
Overview of the model SOLVE. A) dataset curation process, B) feature extraction process, C) model training and prediction recording process and (D) shapley analysis for feature interpretability.

## 3. Materials and Methodology

### 3.1 Dataset Preparation

Previous studies have compiled datasets for enzymes and non-enzymes, but the dataset preparation process has some drawbacks. These ambiguities include the following:

A. Prior studies have limited protein sequence lengths to between 50 and 1000 amino acids. While it is documented that feature extraction and training of ML models on protein sequences longer than 1000 amino acids are computationally intensive, excluding these sequences omits a significant portion of protein sequences present in the UniProt database.
B. As noted earlier, datasets created before 2018 do not include translocase enzymes. ^46^

This study aims to address these challenges in constructing enzyme and non-enzyme datasets. We utilized the publicly available UniProt database^7^ to assemble the datasets (Figure 1(A)). Only reviewed sequences were included, as these are manually annotated and considered superior quality due to extensive curation, including experimental validation. To prepare the enzyme dataset, we have first web-scraped all protein sequences containing an EC number from the UniProt database. For the non-enzyme dataset, we searched for protein sequences in UniProt that lack both EC numbers and catalytic activity. Then, we removed all protein sequences containing unnatural amino acids (e.g., X, J, B) from this filtered dataset to weed out unwanted noises and restricted sequence lengths between 50 and 5000 amino acids. Our final dataset for enzymes and non-enzymes comprised 231907 enzyme sequences and 232474 non-enzyme sequences, and we refer to this dataset throughout the manuscript as Dataset 1. It should be noted, we have collected all the sequences from Uniprot upto 2023 release, which is the most updated dataset for enzyme function prediction till now. Next, for multiple prediction tasks within the enzyme category, we have removed the sequences which have incomplete EC numbers to include only high quality data and constructed subsequent datasets from this primary dataset. To access the effectiveness of SOLVE subsequently we deployed the CD-HIT^53^ program to ensure that the sequence similarity among the gathered sequences, both for enzymes and non-enzymes, remained at the 50%. This implies training set and testing set share less than 50% sequence similarity represent a challenge to any ml models. In this, dataset there are 74764 enzymes and 138256 non-enzyme sequences which is referred as Dataset 2. Throughout this manuscripts, we have evaluated performance of SOLVE in this Dataset-2 in various enzyme hierarchy levels.

### 3.2 Feature Extraction

In the literature, protein’s raw sequences have been one-hot encoded ^48,50^ to extract features and feed them into algorithms to classify them. However, unlike protein structure prediction studies, the sliding window technique ^54^ has received much less attention in enzyme classification. Recently many language models gained much attention to extract meaningful representation from sequence information. ^55–58^ Inspired by these methods, in this study we have used a fine-tuned subsequence tokenization strategy to extract functional features from sequences, which was previously used in other bioinformatics problems^59^; however, this strategy has not yet been used in enzyme prediction problems. Given a protein sequence, a window of a specified size (k) slides over the sequence, one position at a time, extracting k-length overlapping subsequences (k-mers). Feature extraction process has been detailed in Figure 1(B).

Let *S* = *s*_1_*s*_2_*s*_3_…*s_n_* be a protein sequence of *n*, where *s_i_* represents the *i*-th amino acid in the sequence. We define *k* as the length of the k-mer subsequences. The sliding window technique involves extracting subsequences of length *k* from *S*, starting from each position *i* where 1 ≤ i ≤ n − k + 1. Mathematically, the *ii*-th k-mer subsequence can be represented as

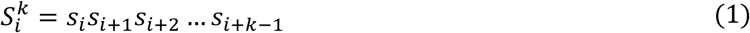

For each starting position *i*, the subsequence 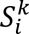 is generated by taking the amino acids from position *i* to *i* + *k* − 1 and the process continues until *i* = *n*– *k* + 1. The method can be represented mathematically as:

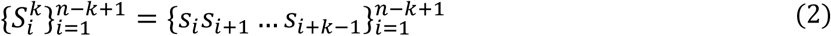

Once all k-mers have been generated, we proceed to tokenize each k-mer. Let *J* be the tokenizer function that maps each k-mer to a unique token:

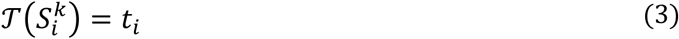

where *t*_*i*_ is the token assigned to the k-mer 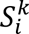.We utilize the tokenizer *J* to encode all k-meric subsequences in the dataset. Considering *E* be the encoding function that uses the tokenizer to encode each k-mer:

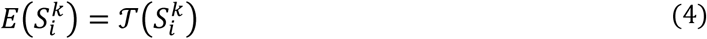

Applying this encoding to all k-mers in the dataset:

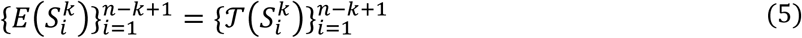

Once all possible k-mers have been assigned a numerical token, the features of every protein sequence are fed into the model along with their corresponding labels. This procedure not only automates feature extraction but also addresses the issue of dimensional non-uniformity and improves memory efficiency. By leveraging the sliding window technique for k-mer generation and subsequent tokenization, we aim to capture local sequence patterns crucial for enzyme classification, thereby improving the predictive performance of our model.

### 3.3 ML model selection and evaluation indices

We have used five different ML models in this study, namely, RF, LGBM, KNN, DT, and SOLVE, to evaluate their performances in different enzyme hierarchy labels. For all the predictions, we used evaluation indices precision, recall, F1 score, sensitivity, and specificity to evaluate the performance of the proposed method. For multi-functional enzyme prediction, we have used slightly different evaluation parameters: subset accuracy, micro F1, macro F1, micro precision, and macro precision.^60^ Subset accuracy is the most appropriate indicator to evaluate multi-label classification. All the mathematical expressions for each evaluation indicator and o the tuned hyper-parameters for each model are given in S1-S2. As enzyme function at L4 label faces significant class imbalance and some classes are very rare, we implement a focal loss inspired penalty into the SOLVE training framework, which adjusts the importance of classes during training to prioritize more on the difficult cases. ^61^ Unlike traditional ml models with default hyper-parameters this approach updates the weight given to different classes dynamically throughout the training phase and boosts the prediction performance overall. The details of focal loss penalty are given in note S3-S4. We analyzed prediction probabilities from SOLVE to understand their correlation with prediction accuracy score. This provides researchers across disciplines access the reliability of SOLVE’s prediction. The dependencies of prediction accuracy with confidence probability are given in fig S4.

## 4. Results and Discussions

### 4.1 Most Discriminative K-mer feature

To optimize the performance of our model, we experimented with k-mer values ranging from 2 to 6 and evaluated the model’s accuracy at each level of the hierarchy. Through systematic analysis, we found that 6-mers consistently yielded the best results across all levels. The box-plot of accuracy scores presented in Figure 2(A) shows K-mer=6 provides the best median accuracy scores consistently for enzyme versus non-enzyme prediction among all the other K-mer values. 6-mer feature descriptor records the highest precision, recall and F1-score of 0.96, 0.92 and 0.94 respectively as it is presented in Figure 2(B). We have also presented ROC curve for different feature descriptors in Figure 2(C). As it is depicted, 6-mer holds the highest AUC score of 0.97, whereas other AUC score of other feature descriptors are almost 22% to 24% lesser. However, using 7-mers overloaded the memory, making it impossible to test beyond this value with our available resources. These results indicate that local sequence patterns can be optimally captured using 6-mers, balancing computational efficiency and predictive performance. In section **4.2**, we have shown the performance of different ML models in all the enzyme hierarchy levels using these 6-mer feature descriptors.

**Figure 2:**
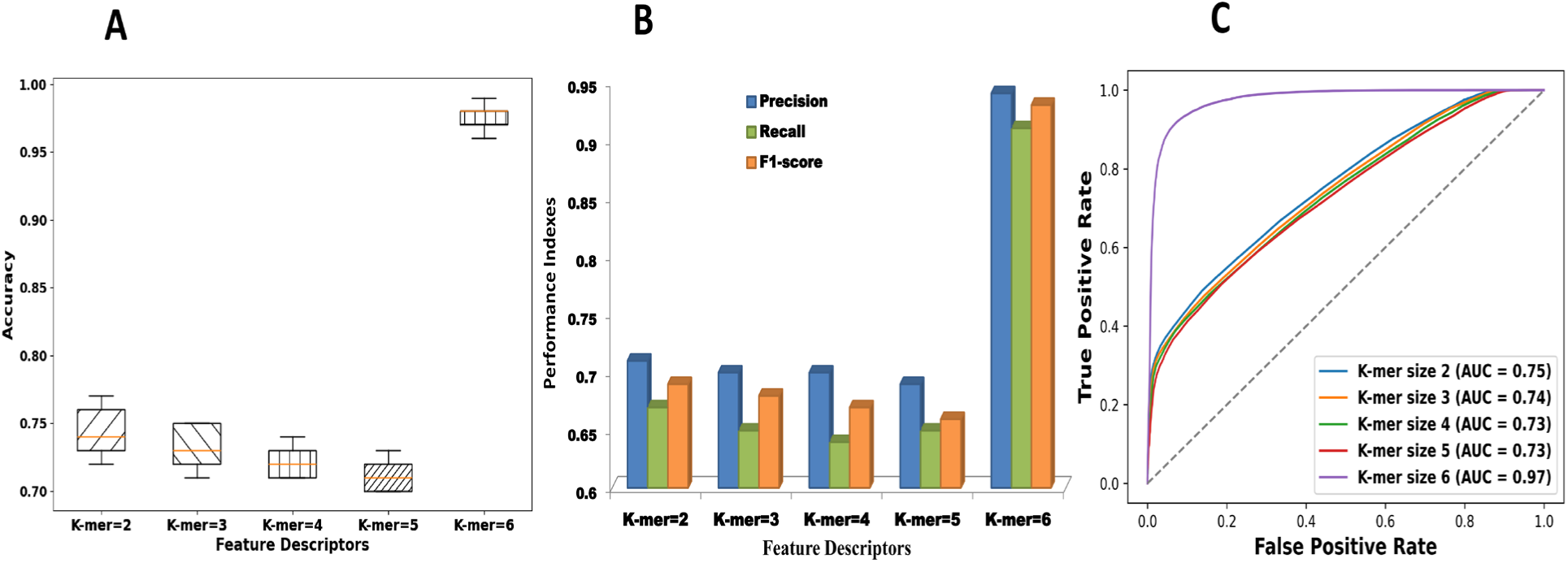
Change of prediction performance with change in k-mer length. (A) The accuracy score of different K-mer feature descriptors are presented as box-plot where orange color horizontal line represents the median accuracy score. (B) Precision, Recall and F1-score of K-mer features and (C) ROC curve for different feature descriptors. The black dotted line represents auc score of a random classifier.

### 4.2 Prediction performance in different levels of enzyme hierarchy

#### 4.2.1 Prediction performance in enzyme vs non-enzyme

To check the performance of our method, we first took Dataset 2 of enzymes and non-enzymes used for training and testing our method. We trained our ML models using 80% random dataset samples and tested their performance on the remaining 20% random samples. To remove any biases present during the prediction, we split the dataset into 80:20 training as well as testing ten times and reported average performance indexes for each prediction task. Multiple ML models, as detailed in the Methods section, were employed to determine the model that exhibits superior performance in distinguishing enzymes from non-enzymes. The performance of these models is presented in Figure 3(A). The figure illustrates that the RF and LightGBM models demonstrated superior performance compared to the DT and KNN models used in this study. Notably, the prediction performance further improved when we employed SOLVE, which combines the predictions of both RF and LightGBM models. This ensemble classifier reached the highest performance in predicting enzymes, with precision, recall, and F1-scores of 0.98, 0.96, and 0.97, respectively. Additionally, the ROC curve shown in Figure S1 coherently demonstrates that SOLVE provides the highest accuracy compared to the other models. It attained 0.98 accuracy in predicting enzymes and non-enzymes, surpassing its closest competitor, the RF model, which achieved 0.97 accuracy.

**Figure 3:**
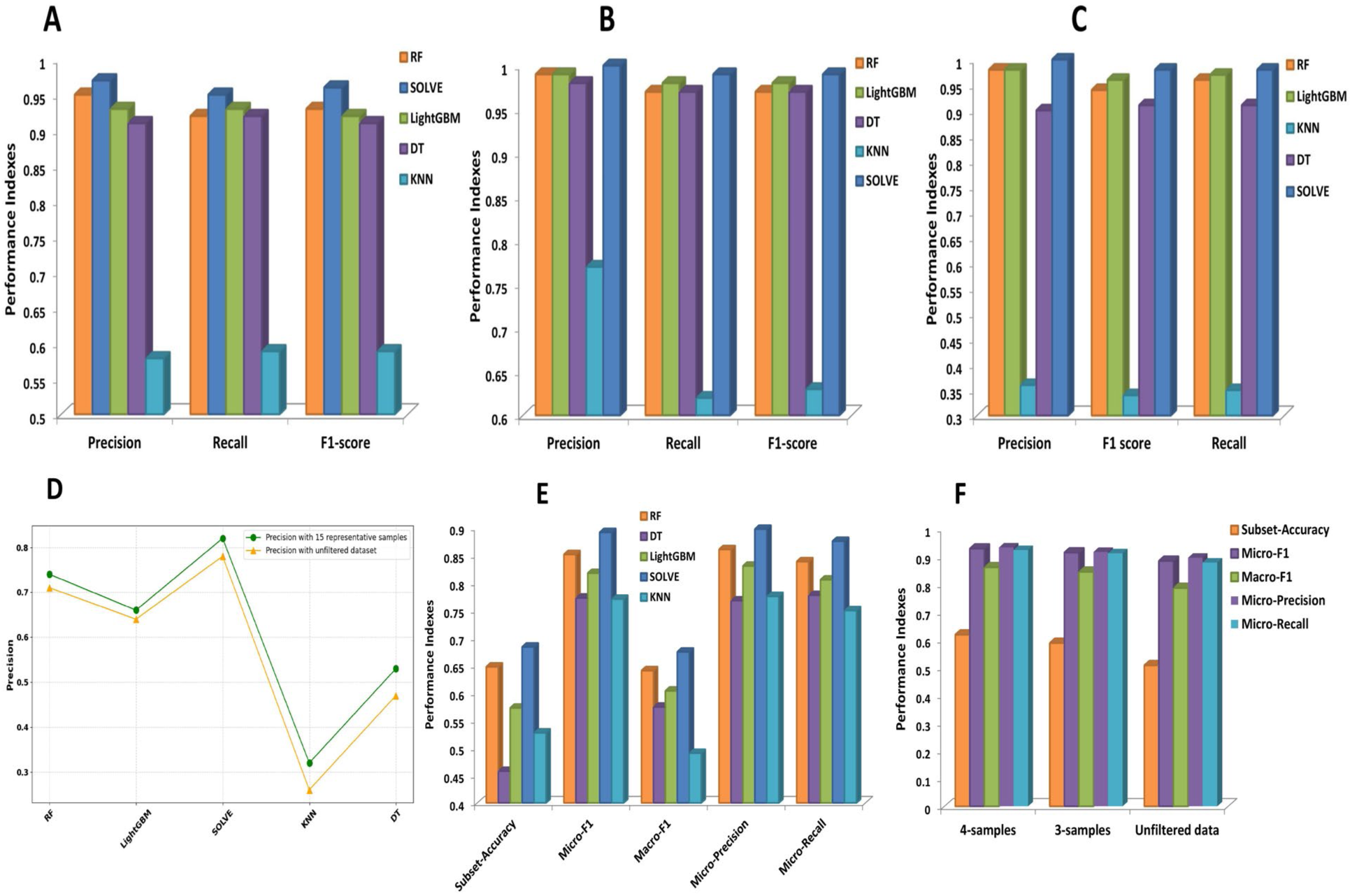
Prediction performance of SOLVE in test-dataset where each sequence share less than 50% sequence similarity.(A) Prediction performance of different ml models in predicting enzymes versus non-enzymes,(B) Performance of five ml models to predict mono and multi-functional enzymes, (C) Prediction performance of five ML models in predicting enzymes main class, (D) Precision scores of five ML models in predicting enzyme substrate class with 15 representative samples per class and in the unfiltered dataset where even less than five samples for some classes are present, (E) Performance of five different ML models used in this study in multi-label multi-functional activity prediction and (F) Performance of different ML models in different minimum distributions of class representatives present in the dataset.

#### 4.2.2. Prediction performance in multi-functional enzymes vs mono-functional enzymes

Next, we advanced to the subsequent level of analysis to determine whether a particular enzyme sequence exhibits mono-functional or multi-functional enzyme activity. Protein sequences with a single EC number were classified as mono-functional enzymes, while those with multiple EC numbers were designated as promiscuous enzymes within the overall enzyme dataset. We evaluated the performance of various ML models on the test dataset using the same approach as previously described. The performance of these models in predicting mono-functional and multi-functional enzymes is presented in Figure 3(B). As Figure 2(C) indicates, SOLVE again outperformed the other ML models in predicting mono-functional and multi-functional enzymes. It reached accuracy, precision, and F1-scores of 0.99, 0.99, and 1.00, respectively, notably superior to the other ML models used in this study. The confusion matrix shown in fig S2(A), reveals that the model correctly predicted 13,958 mono-functional enzymes and 997 multi-functional enzymes. The model misclassified only 1 sample as a multi-functional enzyme and 4 samples as mono-functional enzymes. This result further underscores the model’s efficacy in predicting mono-functional and multi-functional enzymes.

#### 4.2.3 Annotation of Monofunctional Enzymes at *L1* and *L4*

We then moved forward with predicting *L1*, the primary enzyme class, for mono-functional enzymes. To achieve this, we excluded all multi-functional proteins from the enzyme dataset and removed enzyme sequences with incomplete EC numbers at *L4*. This process resulted in 54,232 samples belonging to one of the seven different enzyme classes. At this level, we found weights of 2:1.5:0.2 for the RF, LightGBM, and KNN models in the ensemble learning framework, demonstrating the best performance. The performance of other weight ratios is systematically presented in table S4. As shown in Fig 3(C), SOLVE delivered the best performance predicting the main enzyme class, achieving a precision, F1 score, and recall of 1.00, 0.99, and 0.99, respectively. The RF and LightGBM models also performed very well, although their performance was slightly inferior to SOLVE’s. The confusion matrix shown in Figure S2(B) illustrates that the true positive predictions for each enzyme class are significantly higher with SOLVE, further validating the superiority of this model for enzyme function prediction.

The results of predicting enzyme substrate classes, i.e., *L4* level, are presented in Fig. 3(D). It is important to note that predicting enzyme function at *L4* is particularly challenging due to the extreme sparsity of representative samples for each substrate class. In some cases, only one or two samples are available for a given substrate class in the entire dataset. We first filtered the dataset to include only those substrate classes with at least 20 samples, removing other classes. As shown in Fig.3(D), to our surprise, RF almost performed at par with SOLVE. We also evaluated our method’s performance on the unfiltered dataset. As depicted in Fig.3(D), even in the unfiltered dataset—where many substrate classes have very few samples—SOLVE achieves remarkable performance with a precision of 0.77. Additionally, we have conducted another analysis to observe how the performance at the *L4* level increases and reaches a plateau when we take top K accuracy predictions other than a single prediction. The result in Fig. S3 clearly shows that given a considerable number of enzyme classes at *L4,* if one considers only the top 12 predictions, the accuracy goes above 90%. This suggests that the prediction results presented in this study are not arbitrary, showing a strong correlation between the enzyme function and protein subsequence patterns. Experimentally, it is also helpful to identify the likely function of an uncharacterized enzyme within the top 10 or 20 predictions, helping to narrow down the vast range of possible enzyme functions. Hereby, we emphasize that there is room for significant improvement in substrate class prediction, and increasing the number of samples for rare substrate classes is crucial for training more effective ML models.

#### 4.2.4 Annotation of Multi-label Multi-functional Enzymes at *L1* and *L4*

First, we excluded multi-functional enzyme sequences with incomplete EC numbers at the *L4* level. It is important to note that an enzyme can exhibit promiscuous activity at any level of the enzyme hierarchy, from *L1* to *L4* of the EC number. Since *L1* represents the main enzyme class, we initially predicted the multi-functional enzyme activity based on the primary enzyme function, and the results are presented in Table S3. The RF and LightGBM models demonstrated vital predictive metrics as tabulated, while the KNN and DT models performed inferiorly. Notably, when applying a soft voting ensemble with weights of 10:5:1 for RF, LightGBM, and DT, respectively, SOLVE outperformed others in predicting multi-functional enzymes at *L1;*other weight ratios with their performance are presented in Table S1.It attained a subset accuracy of 0.65, a micro F1-score of 0.97, and a micro precision of 0.97 (see Table S1). Following the successful prediction of multi-functional enzymes at *L1*, we predicted multi-functional enzyme activity down to *L4*, corresponding to the substrate-binding class. We emphasize that predictions at this level are more challenging due to the dataset’s many unique class combinations and the meagre sample size for each unique substrate class. Consequently, few studies have attempted to predict multi-functional enzyme classes down to the last label.

Initially, we filtered our dataset to ensure at least five samples for each EC number present, a procedure commonly applied in other studies in this domain. This filtering step ensures that the ML models have atleast some samples for each class combination during training. The results of multi-label, multi-functional enzyme prediction down to *L4* are presented in Fig.3(E). SOLVE reached a subset accuracy of 0.68, approximately 2% and 11% better than the RF and LGBM models, respectively. Additionally, SOLVE provided a micro F1-score of 0.89 and a micro precision of 0.90, outperforming the other models used in this study. Through rigorous testing with SOLVE, we found that a combination of RF, LGBM, and DT with an optimal ratio of 4:5:0.25 yields the best performance. The results for various other weighting ratios are provided in table S2.

We also tested the SOLVE performance under different data distribution scenarios to address the limitations of having only five representative samples per class. As shown in Fig.3(F), the performance metrics of SOLVE decrease when the minimum number of samples per class is gradually reduced. The subset accuracy drops from 0.68 to 0.62 when the minimum number of samples per class decreases from five to four. We also evaluated the model’s performance on the unfiltered dataset, as represented in Fig.3(F). In this scenario, many new class combinations that the model did not encounter during the training phase were present, making it challenging for the ML model to predict the multiple functions of these enzymes. Interestingly, even in this situation, SOLVE reached a subset accuracy of 0.51, a micro F1-score of 0.89, and a micro precision of 0.90. These results suggest that our model effectively recognizes the underlying patterns in enzyme sequences to predict specific catalytic activities, even with limited data.

#### 4.2.6 Benchmarking SOLVE on independent dataset to compare the performance among different methods

In the Dataset preparation section, we have already mentioned that we have trained our models with all the UniProt sequences until 2023. We have curated a dataset of enzyme and non-enzyme sequences from the reviewed portion of Uniprot database which came after 2023 as these sequences are not part of the SOLVE model development. For curating this dataset, we have followed the same criteria as we have mentioned in the Dataset preparation. We have collected 505 enzyme and 694 non-enzyme sequences and tested the performance of SOLVE along with state-of-the-art methods. The figure presented in 4(A) clearly demonstrates the superiority of SOLVE in discriminating enzymes from the non-enzymes ones. SOLVE has achieved precision and F1-score of 0.84 and 0.82, respectively. SOLVE has achieved 9% and 8% improvements in precision and F1-score over the second best model ECPred in the independent dataset. Moreover, it is important to note that CLEAN cannot discriminate enzymes from the non-enzyme ones. As a result when presented an unknown non-enzyme sequence, CLEAN will still assign an EC number, which can lead to misannotations in high throughout protein function prediction. SOLVE has also achieved improvements in precision and recall of 17% and 18% in comparison with deep learning model, DeepEC. Then, we tested different methods’ performance on level 1 and level 4 in this Uniprot 2024 dataset consisting of 505 enzymes for which EC numbers upto 4^th^ digit are known. The results shown in Figure 6(B) depicted that SOLVE gave a slightly greater performance compared to CLEAN in enzyme main class prediction. SOLVE achieves accuracy and F1-score of 0.80 and 0.76, whereas CLEAN delivers accuracy and F1-score of 0.76 and 0.74, respectively. DeepECtransformer and DeepEC achieve accuracy of 0.67 and 0.58, respectively. SOLVE achieved improvements in F1-score of 4%,12% and 20% compared with DeepECtransformer, Proteinfer and DeepEC respectively. Then we moved on to evaluate performance of SOLVE with contemporary methods in the L4 level. We have found that in this level of enzyme hierarchy, using combinations of 6-mer and 4-mer feature descriptors delivers better performance. So in the L4 level we have presented the performance of SOLVE with combinations of K-mer features along with other tools for enzyme EC number annotations in figure 4(C). The detailed ablation study of how different combination of feature descriptors influencing the prediction results is presented in Figure S4. SOLVE once again provides better performance in the *L4* label than CLEAN and other four methods. SOLVE has achieved accuracy and F1-score of 0.48 and 0.36, which are 3% and 5% improvements over CLEAN. SOLVE also delivers 11% and 14% better F1-score compared to recently developed tools DeepECtransformer and Proteinfer. We have also curated a multi-functional enzyme sequences from Uniprot 2024 release and evaluated the SOLVE’s performance with the other tools. The figure presented in 4(D) clearly demonstrates that SOLVE outperformed all other ec annotation tools for enzyme promiscuity predictions. SOLVE provides precision and f1-score of 0.89 and 0.73 respectively which are almost 42% and 36% improvements over the second-best model CLEAN. We then evaluated the SOLVE’s performance along with other tools in the dataset where testing enzyme sequences share less than 40% sequence similarity with the training set ones. In such a low sequence similarity scenario SOLVE delivers precision and F1-score of 0.8 and 0.78 respectively as it is depicted in figure 4(E), whereas CLEAN achieved precision and F1-score of 0.77 and 0.75 respectively. We have used another independent test dataset, New-392, which contains a variety of enzymes with different EC numbers. It is important to note that this dataset was constructed from the SwissProt after the 2022 release, and we have trained our model with the Uniprot dataset till 2023.. We first removed the sequences already present in the training set and made predictions for the remaining ones. We have compared our results with those of DeepECtransformer^51^, ProteInfer^48^, DeepEC^50^, and ECPred^47^ for benchmarking. We have found that SOLVE outperformed all of these methods in performance indicators. SOLVE’s precision and recall score is about 36% and 26% higher than ECPred and about 4% and 3% higher than DeepECtransformer. This demonstrates that our method, SOLVE, is more effective in predicting enzyme functions than these existing tools. We also compared SOLVE’s performance with the CLEAN model in New-392, which shows that SOLVE underperforms slightly compared to CLEAN. CLEAN achieved only 4% greater F1-score than SOLVE. Moreover, we have taken four bacterial proteins from the NCBI database and tested our model on them. SOLVE correctly characterized the enzyme class of 4 out of 5 sequences, demonstrating its efficacy on unseen sequences. Interestingly, CLEAN can correctly predict the true enzyme class for only one enzyme sequence. The details of the predictions are given in note S3 (Table S5). This stresses that effectiveness of SOLVE in predicting enzyme functions compared with CLEAN.

**Figure 4:**
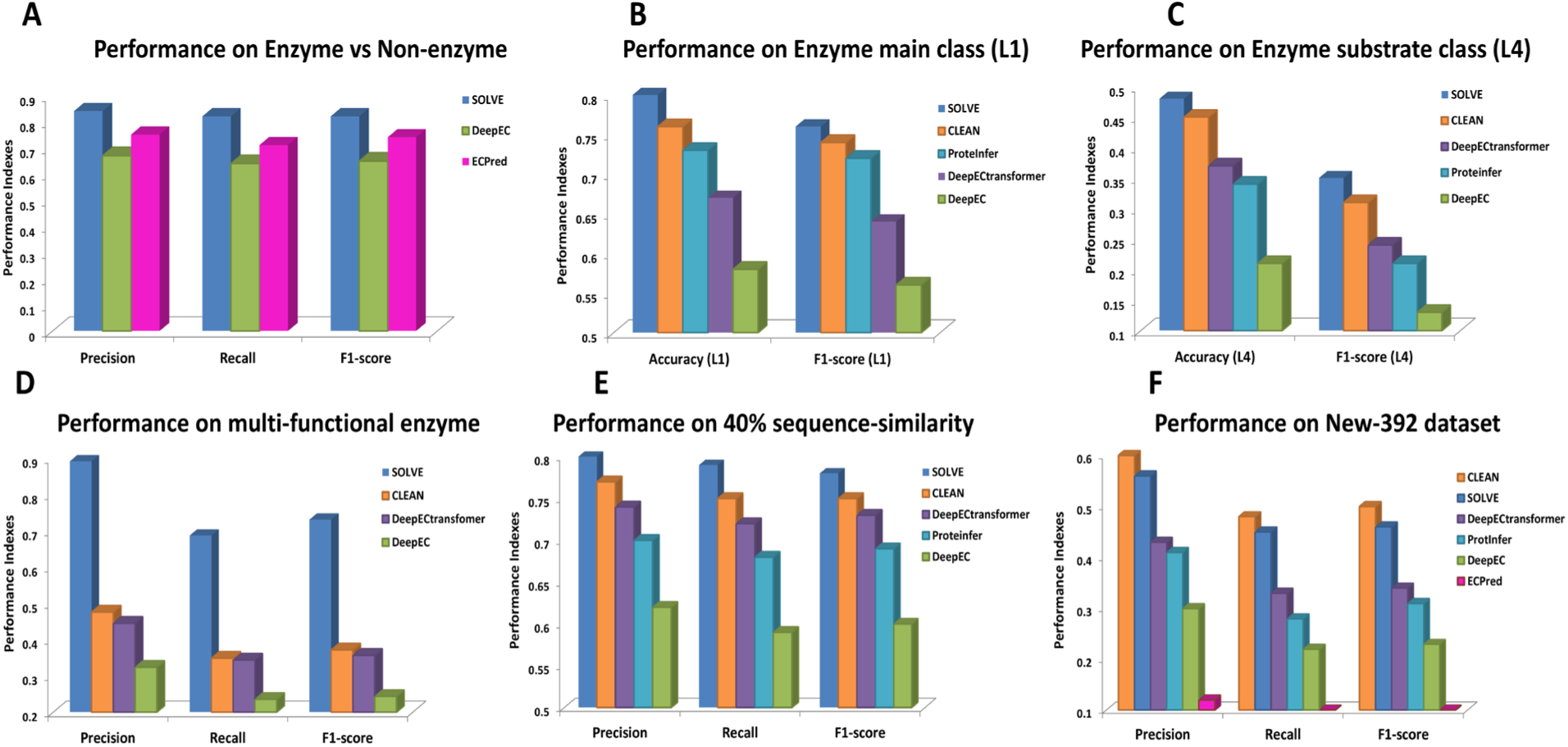
Comparison of SOLVE with the state-of-the-art models for enzyme function prediction. (A) Prediction performance of SOLVE for for predicting Enzymes vs Non-enzymes in Uniprot 2024 dataset. Three methods, CLEAN, DeepEC and ECPred are taken for comparison, (B) Prediction performance of SOLVE, CLEAN, DeepECtransformer, DeepEC in prediction of enzyme main class on UniProt 2024 dataset, (C) Prediction performance of SOLVE in *L4* label of UniProt 2024 dataset compared with SOLVE, DeepECtransformer and DeepEC and (D) Prediction performance of SOLVE in Uniprot 2024 multi-functional datset with three contemporary methods, (E) Evaluation of SOLVE’s performance on 40% sequence similarity dataset and (F) Prediction performance of different methods in New-392 dataset for enzyme prediction at *L4* label.

#### 4.2.7 Feature importance analysis

Feature importance analyses are crucial as they provide insight into which features most influence model predictions, enabling better interpretability and trust in ML models. Many deep learning models lack interpretability, which in turn creates obstacles to understanding the actual biological phenomena that guide ML models’ prediction, such as the CLEAN method. Understanding feature importance can significantly enhance model refinement and optimization by focusing on the most predictive variables, improving overall model accuracy and robustness. Over the last decade, many interpretable AI models have been developed to explain the ML model’s outcomes.^62,63^ This study focused on interpreting our model by calculating their average feature importance with Shapley analyses, which provides critical insights about the underlying mechanism of the black-box ML model’s decision process and is widely used in different domain problems.^59,64–66^ We selected two proteins from PDB (IDs: 5MO4 and 6OIM) and excluded these sequences from the training dataset. Our model accurately identified these enzyme’s EC numbers.

We computed the feature importance for each 6-mer amino acid subsequence and then highlighted the top 30 subsequence stretches that contributed most significantly to the model’s predictions within the protein’s tertiary structure (Figure 5). Annotating these stretches on the protein structures revealed that our model effectively captures critical biological patterns within the protein’s primary sequence. These top-performing features are located within these proteins’ orthosteric binding sites and allosteric sites, as illustrated in the Figure 5. In recent times, numerous experimental and computational studies have focused on identifying allosteric site and investigating the mechanism of signal transduction within proteins.^67–71^ ABL1 kinase is extremely important in signal transduction and often targeted as cancer therapies particularly for chronic myeloid leukemia (CML).^72,73^ For the ABL1 kinase, asciminib drug binds to the myristoil pocket which is also the allosteric site of this enzyme.^74,75^ In contrast, nilotinib drug binds to the ATP-binding pocket of this enzyme. For, G12C mutant kras enzyme is an important target in cancer research.^76^ GDP binds to orthosteric site of this kras enzyme and a covalent inhibitor AMG 510 binds to the allosteric site.^77,78^ The top 15 features are predominantly found within or adjacent to the allosteric site regions, while the subsequent top 15 features correspond to the orthosteric binding sites for both proteins.

**Figure 5:**
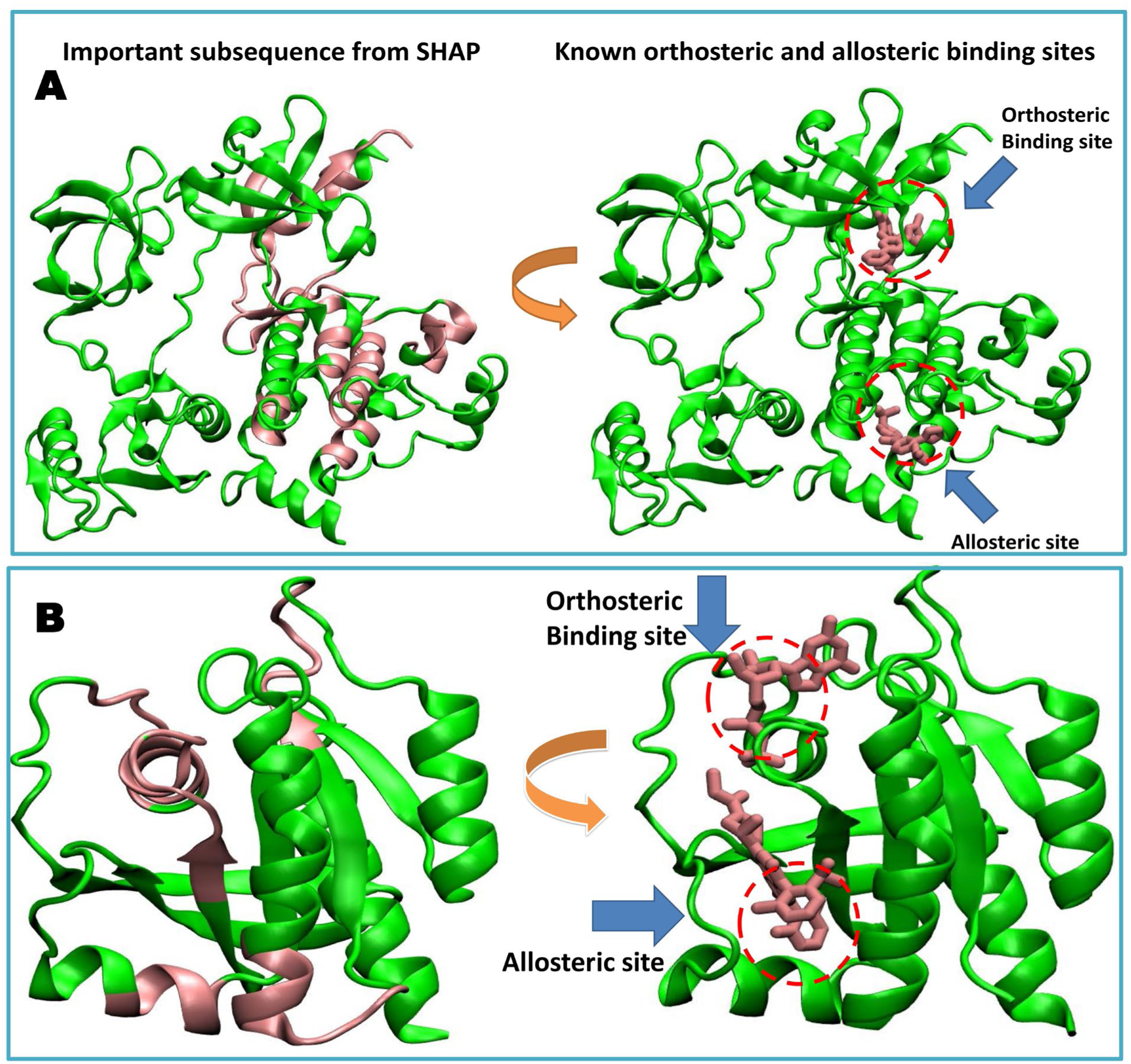
Important 6-mers extracted from feature importance analysis are mapped onto the 3D structure of two enzymes. (A) PDB ID: 4Q21 (kras protein) and (B) PDB ID: 5MO4 (ABL kinase protein). The top thirty important 6-mers calculated from SHAP analysis are shown in the left panel, and binding and allosteric sites are marked in red circles. The two proteins’ actual binding and allosteric regions are shown in the right panel.

## 5. Conclusions

Prediction of enzyme function is a fundamental challenge in biology. Due to the time-consuming nature of experimental methods, new computational tools based on biochemical features, sequence similarity, and ML-based methods are necessary for high-throughput screening of enzyme functions. However, most of the previous enzyme annotation models needed improvements in feature extraction, interpretability, and their ability to adapt to unseen datasets. Most of them struggle to distinguish between enzyme and non-enzyme sequences, leading to misclassification and limited generalization.

In this work, we developed SOLVE, a supervised ensemble learning framework to predict enzyme functions. This ensemble model leverages numerical tokenization of raw protein sequences, allowing it to capture key sequence patterns without relying on predefined biochemical features. This feature extraction strategy based on NLP provides a route to predict enzyme functions based on their short subsequence pattern. SOLVE distinguishes enzymes from non-enzyme sequences, predicts mono- and multifunctional enzymes, and provides substrate-binding classifications down to the substrate-binding level with greater accuracy. SOLVE offers valuable insights into enzyme function and activity by pinpointing essential functional regions, such as those involved in oncogenic proteins like KRAS. Our model’s ability to directly link sequence patterns to enzymatic activity provides deeper insights into protein’s functionality. These improvements may position SOLVE as a critical tool in accelerating drug discovery, genomics research, enzyme engineering and the functional annotation of novel enzymes. Moreover, this framework is not only limited to enzymatic activity prediction, but this method can be used to predict other functional proteins. User-friendly framework of SOLVE enables broader adaptation and streamlined enzyme EC number prediction by the research community. However, we acknowledge that there is room for improvement for enzyme function in the *L4* level. Moreover, for feature extraction, incorporating protein structural features and information about the catalytic site of enzymes may significantly enhance the performance of this model. These points will be the main focus of our future work.

## Data and Code Availability

All the data supporting supporting the findings are included in the manuscripts. Additionally, the datasets, all the codes and trained models can be accessible from Github via the following link: (https://github.com/saikat-ai/Enzyme_prediction).

## ACKNOWLEDGMENTS

This research is supported by the Department of Science and Technology SERB grant CRG/2020/000756 for funding. All the authors acknowledge IACS for super-computing facilities. S.D. acknowledges IACS for financial support.

## Notes

Authors declare no conflict of interest.

