## Supporting Information for "Prediction of Enzyme function using interpretable optimized Ensemble learning framework"

**The SI file includes:**

SupplementaryText
Fig. S1 to S5
Table S1 to S5

### Supplementary text

#### Note 1. Mathematical equations for various evaluation indices

$$\text{Precision} = \frac{\text{TP}}{\text{TP} + \text{FP}} \tag{6}$$

$$\text{Sensitivity} = \frac{\text{TP}}{\text{TP} + \text{FN}} \tag{7}$$

$$\text{F1 Score} = 2 \times \frac{\text{Precision} \times \text{Recall}}{\text{Precision} + \text{Recall}} \tag{8}$$

$$\text{Specificity} = \frac{\text{TN}}{\text{TN} + \text{FP}} \tag{9}$$

$$\text{Subset Accuracy} = \frac{\text{Number of correctly predicted subsets}}{\text{Total number of subsets}} \tag{10}$$

$$\text{Micro F1} = \frac{2 \times \text{TP}_{\text{micro}}}{2 \times \text{TP}_{\text{micro}} + \text{FP}_{\text{micro}} + \text{FN}_{\text{micro}}} \tag{11}$$

$$\text{Macro F1} = \frac{1}{N}\sum_{i=1}^{N} \text{F1}_i \tag{12}$$

$$\text{Micro Precision} = \frac{\sum_{i=1}^{N} \text{TP}_i}{\sum_{i=1}^{N} (\text{TP}_i + \text{FP}_i)} \tag{13}$$

$$\text{Macro Precision} = \frac{1}{N}\sum_{i=1}^{N} \frac{\text{TP}_i}{\text{TP}_i + \text{FP}_i} \tag{14}$$

TP, TN, FP, and FN represent true positive (correctly predicted enzyme), true negative (correctly predicted non-enzyme), false positive (incorrectly predicted non-enzyme as an enzyme), and false negative (incorrectly predicted enzyme as non-enzyme), respectively. Note that equation numbers have been kept in continuation of the equation numbers from the main manuscript.

#### Note 2. Hyper-parameters for deployed ML models

Here, we have delineated the models and hyperparameters we used and refined to build SOLVE.

```
dt
=DecisionTreeClassifier(criterion='gini',max_depth=5,min_samples_split=2,min_samples_
leaf=1,max_features=None)

rf =
RandomForestClassifier(n_estimators=100,criterion='gini',max_depth=10,min_samples_spl
it=2,max_features='sqrt')

lgbm = lgb.LGBMClassifier(num_leaves=31,learning_rate=0.1,max_depth=-
1,n_estimators=100,min_data_in_leaf=20)

knn =
KNeighborsClassifier(n_neighbors=5,weights='uniform',algorithm='auto',p=2,metric='min
kowski')

solve = VotingClassifier(estimators=[('dt', dt), ('rf', rf), ('lgbm',
lgbm)],voting='soft',weights=[weight1, weight2, weight3],n_jobs=-1)
```

### Note 3. Four bacterial protein predictions from the NCBI database

To further validate our model, we subjected SOLVE to predict the enzymatic classes of proteins at *L1* with experimentally verified enzymatic activities that were not part of the training dataset. Specifically, we focused on entries yet to be included in the Swiss-Prot database. We ensured that this small dataset comprised enzymes that only exhibited a single functionality and were not promiscuous. To avoid bias in the dataset, we manually curated a dataset of four random proteins from various bacterial genome databases within NCBI. The predictions and their comparison with experimentally validated classifications are presented in table S5.

As depicted, our model correctly predicted the enzymatic class at *L1* for three out of four entries, demonstrating its reliability in detecting local patterns in enzymes and accurately classifying them. However, SOLVE failed on WP_256936340 protein, which has 27 AA. The inefficiency of our model may be due to the inclusion of sequences shorter than 50 amino acids, as our dataset predominantly consists of proteins with lengths ranging from 50 to 5000 amino acids. These shorter sequences may need more information for accurate classification, potentially affecting the model's overall performance. These results underscore the model's robustness in enzyme classification and highlight areas for further refinement, particularly in handling shorter protein sequences.

### Note 4. Focal loss penalty

$$L_{focal} = -\alpha(1 - p_t)^{\gamma} log\, p_t$$

Here α is the balancing parameter, γ is the focusing parameter, $p_t$ is the predicted probability for the class and $L_{focal}$ is the focal loss function. We adjusts the gamma parameter to tune the weightage of misclassified class in successive rounds of training.

### Note 5. Algorithm of SOLVE with iterative weight tuning

**Step 1**: **Initialize Parameters**

Set the initial sample weights: $w_0$=[1,1,…1]

**Step 2**: **Iterative Training Loop**

For t=0, 1, 2, T-1:

1. Train the model”

   Train SOLVE model $S_t$ on the dataset G with current sample weights $w_t$:

   $$S_t = SOLVE\ (G, w_t)$$

2. Predict class probabilities for each sample i:

   $$P_t(X_i) = [p_{t,1}, p_{t,2}, p_{t,k}],\ i\epsilon\ \{1,2,\ldots N\}$$

3. Clip probabilities of the predicted class:

   Ensure that none of samples gets 0 probability during the training phase

   $$P_t^{clipped}(X_i) = clip\ (P_t\ (X_i),\epsilon,\ 1\text{-}\epsilon)$$

4. **Compute Focal Weights**

   For each sample i, calculate the focal weight using the predicted probability of the true class $y_i$:

   $$W_{t+1,I} = (1\text{-}\ p_t^{clipped}(X_i)[y_i]\ )^{\gamma}$$

5. **Update Sample Weights**

   Update the sample weights for the next iteration:

   $$W_{t+1} = [w_{t+1,1}, w_{t+1,2}, \ldots.. w_{t+1,k}]$$

**Step 3: Final Model**

After T iterations, SOLVE with last set of sample weights $w_T$

$$S_T = SOLVE\ (G, w_T)$$

**Fig. S1**.
**The ROC curve for five different ML models predicting enzymes vs non-enzymes.**

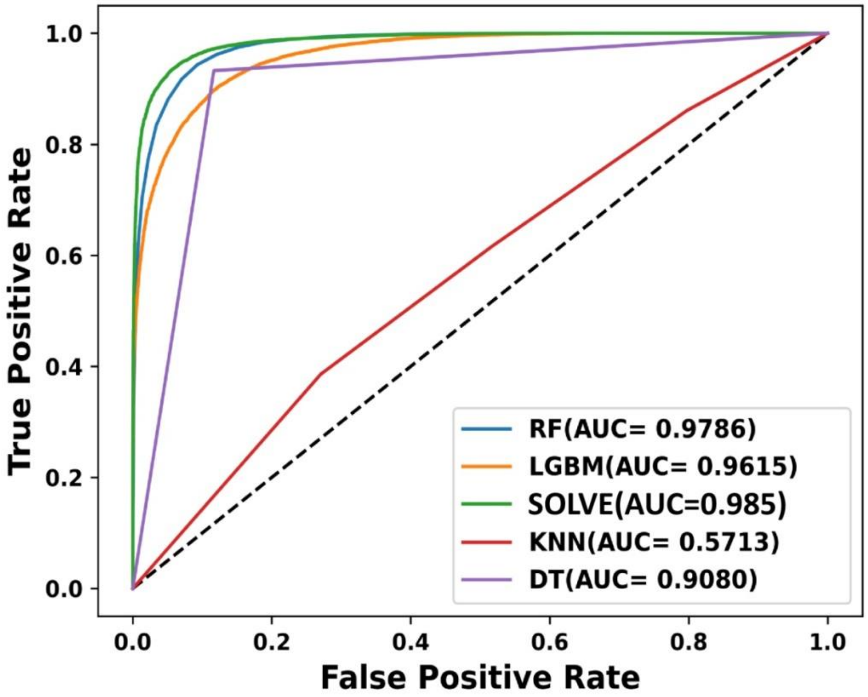

**Fig. S2**.
**Confusion matrices for predictions at different levels.**(**A**)Confusion matrix of SOLVE in predicting mono vs multi-functional enzymes, and (**B**) Confusion matrix of SOLVE in predicting enzyme main class prediction.

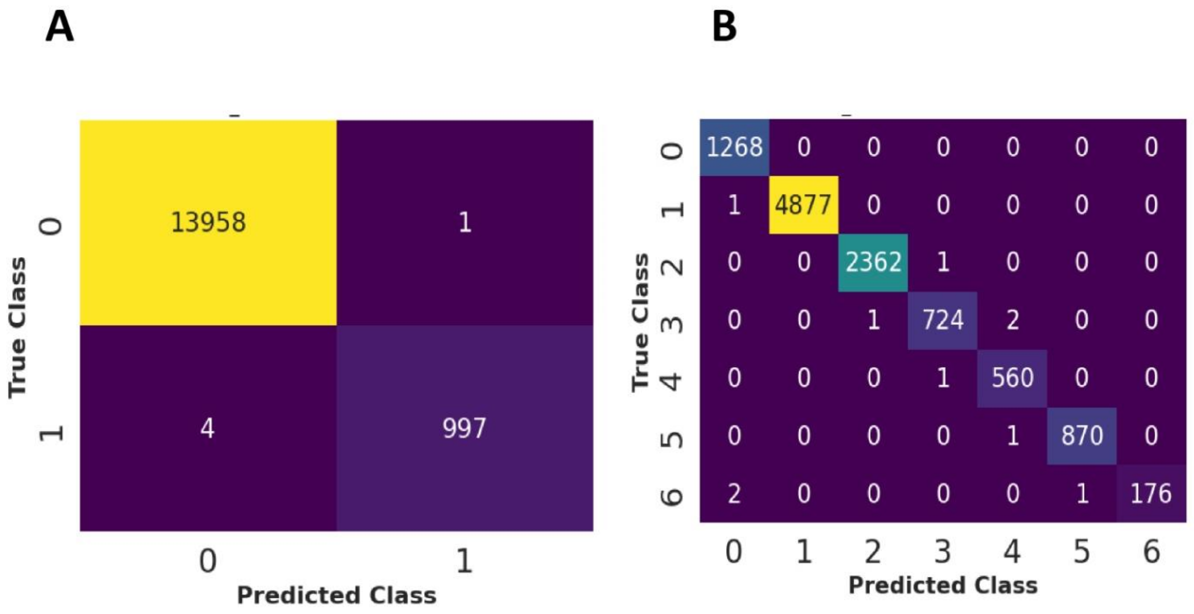

**Fig. S3**.
**Top K accuracy of SOLVE at *L4* level**. The model reaches 0.90 accuracy after 12 predictions and 0.80 after top 2 predictions.

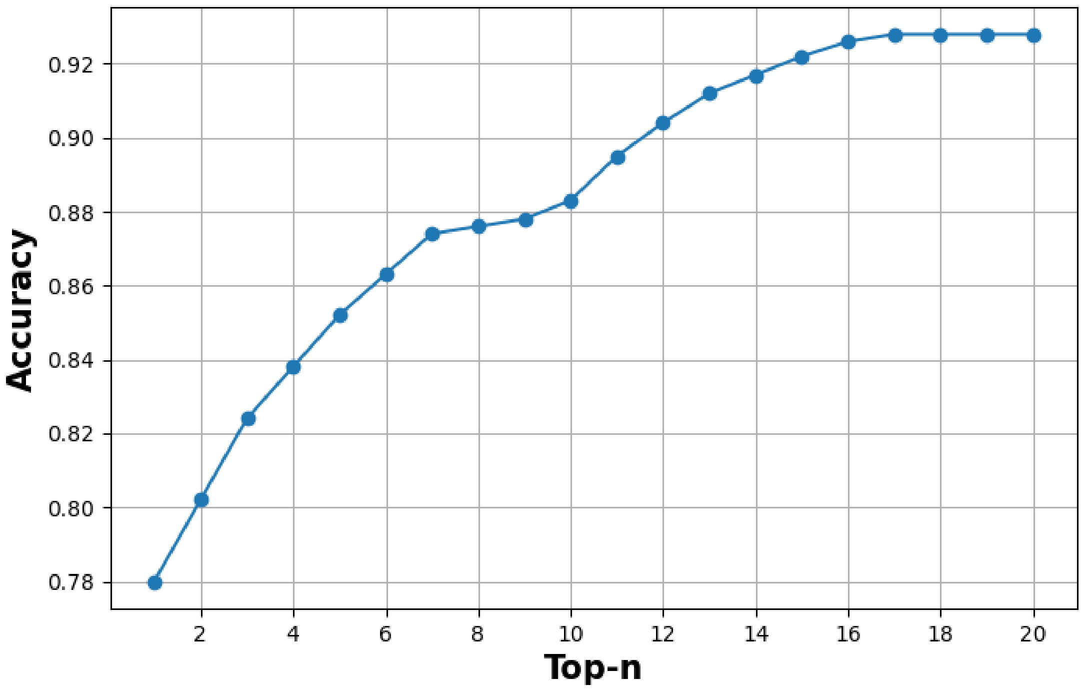

**Fig S4:**

**Confidence score vs Accuracy plot.** When the confidence score is very low 0.1, accuracy is also very low 0.15. As the confidence score of SOLVE increases, accuracy is also increasing. Accuracy score reaches a plateau after confidence score of 0.5.

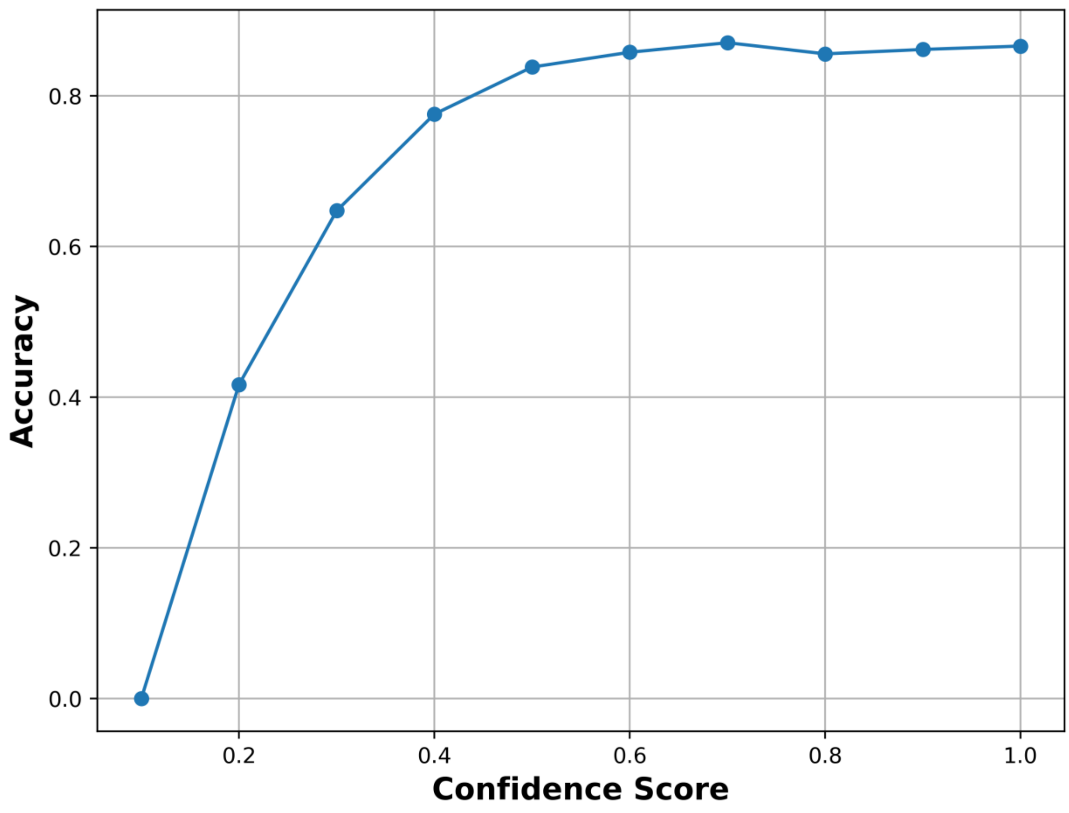

**Fig. S5.**

**Ablation study of how different features influence accuracy at the *L4* label of enzyme prediction.** The combination of 6-mer+4-mer feature descriptor records the best accuracy of 0.48.

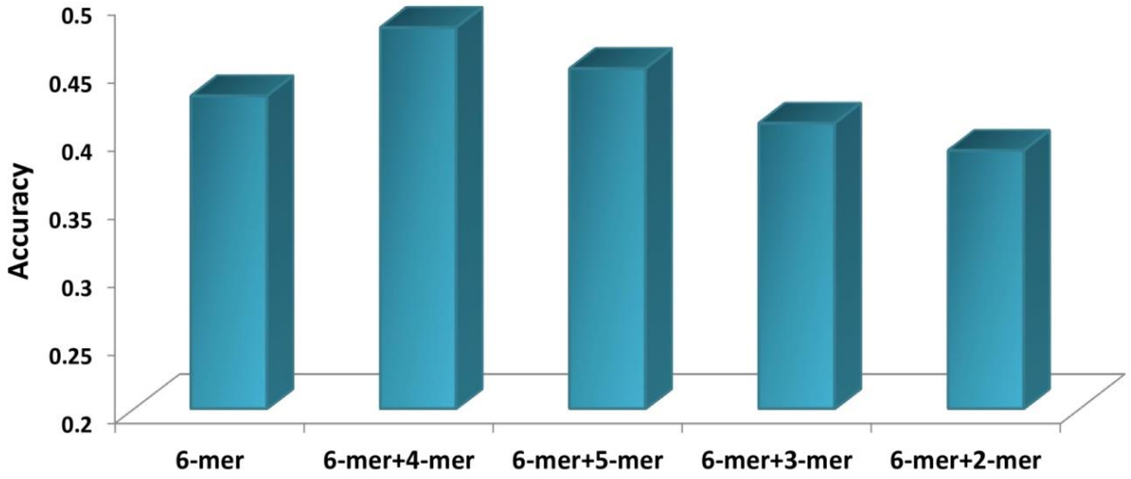

**Table S1**.
**Different weight ratios of SOLVE classifier in predicting *L1* of multi-functional enzyme.**

| Weight Ratio of RF: LGBM: DT | Subset Accuracy |
|---|---|
| 10:5:1 | 0.65 |
| 5:3:1 | 0.52 |
| 8:4:1 | 0.62 |
| 4:5:1 | 0.48 |
| 4:5:6 | 0.32 |

**Table S2.**
**Different weight ratios of SOLVE classifier in predicting *L4* of multi-functional enzyme.**

| **Weight Ratio of RF: LGBM: DT** | **Subset Accuracy** |
|---|---|
| 4:5:0.25 | 0.52 |
| 5:4:0.25 | 0.49 |
| 4:5:2 | 0.45 |
| 4:8:0.25 | 0.51 |

**Table S3**.
**Performance of five different ML models in predicting the *L1* label of the multi-functional enzyme.**

| Model | Subset accuracy | Micro F1 | Macro F1 | Micro Precision | Macro Recall |
|---|---|---|---|---|---|
| RF | 0.602 | 0.963 | 0.882 | 0.963 | 0.963 |
| KNN | 0.507 | 0.935 | 0.770 | 0.928 | 0.941 |
| DT | 0.316 | 0.951 | 0.851 | 0.952 | 0.953 |
| LGBM | 0.512 | 0.961 | 0.874 | 0.965 | 0.958 |
| SOLVE | 0.654 | 0.970 | 0.886 | 0.972 | 0.969 |

**Table S4**.
**Weight ratios of SOLVE classifier in predicting *L1* of mono-functional enzyme.**

| **Weight Ratio of RF: LGBM: DT** | **Accuracy** |
|---|---|
| 2:1.5:0.2 | 0.99 |
| 1.5:2:0.2 | 0.98 |
| 2:1.5:1 | 0.96 |
| 1.5:2:1 | 0.95 |

**Table S5.**
**Experimentally identified enzyme prediction using SOLVE and CLEAN with *L1* label.**

| NCBI Locus | True Class | SOLVE Prediction | CLEAN Prediction |
|---|---|---|---|
| WDD90342 | Isomerase | Isomerase | Oxidoreductase |
| WDD90234 | Translocase | Translocase | Ligase |
| WP_256936340 | Oxidoreductase | Ligase | Hydrolase |
| AKJ08511 | Hydrolase | Hydrolase | Transferase |
| AKJ08508 | Transferase | Transferase | Transferase |
